# Possible adaption of the 2022 Monkeypox virus to the human host through gene duplication and loss

**DOI:** 10.1101/2022.10.21.512875

**Authors:** Annika Brinkmann, Claudia Kohl, Katharina Pape, Daniel Bourquain, Andrea Thürmer, Janine Michel, Lars Schaade, Andreas Nitsche

## Abstract

Poxviruses are known to evolve slower than RNA viruses, with rearrangements such as gene gain and loss as the main driver for host adaption. In 2022 the world is being challenged by the largest global outbreak so far of Monkeypox virus, and the virus seems to have established itself in the human community. Here we report five MPXV genomes with extensive gene duplication and loss, including duplications of up to 18,000 bp to the opposed genome end, and deletions at the site of insertion of up to 16,000 bp, as a possible adaption to the human host.

## Introduction

Monkeypox virus (MPXV) is a species of the Orthopoxviruses (OPV), a genus that includes camelpox (CMPV), cowpox (CPXV), vaccinia (VACV) and variola virus (VARV). Following the eradication of the latter, MPXV was described as the most important OPV species, and in the years following VARV eradication concern was raised that MPXV might fill the vacant epidemiological niche of variola virus [1-3]. However, between the first identified case of human monkeypox in 1970 in the Democratic Republic of the Congo (DRC) and the first outbreak in a non-endemic country with 81 identified cases (in 2003, USA), only few cases of human monkeypox were reported outside the endemic region of DRC [4,5]. In the following years, further outbreaks were reported in the Sudan (2005) and in several countries in Central and West Africa (2017–2018), some of which had not reported any cases of human poxvirus for almost 40 years [6-8]. In 2018 and 2019, cases connected to travelers infected in Nigeria were reported outside of Africa for the second time, raising increased awareness in the UK, Israel and Singapore [9-11]. Not only had monkeypox been “reported from more countries in the past decade than during the previous 40 years” [12], but also human cases had more than doubled in that period. However, up to this point, no more than seven serial human transmission events were reported, and MPXV was considered unlikely to maintain itself in human communities without repeated zoonotic spill-over events from its (yet unknown) natural host [13-18].

In May 2022, cases of monkeypox were reported in the UK, followed by Spain, Portugal, the US and several countries all over the world [19-21]. On 13 October 2022, 72,874 cases were reported in 109 countries, 102 of which had never reported monkeypox before [22]. Total deaths were 28, in Nigeria (7), Brazil (5), Ghana (4), Cameroon (2), Spain (2), United States (2), Belgium (1), Cuba (1), Czechia (1), Ecuador (1), India (1) and Sudan (1) were reported. Phylogenetically, the current outbreak strain belongs to clade II (former West African clade) generally considered to be less virulent than clade I (former Congo Basin clade) [23]. It forms a divergent branch (clade IIb) with 46 nucleotide differences to Nigerian strains from 2019 and 16 nucleotide differences to a strain from a patient travelling from Nigeria to the USA in 2021 [24,25]. The high mutation rate in the current outbreak strain, which was previously estimated as 1–2 nucleotides/genome/year for poxviruses and the distinct signature of nucleotide changes (GA>AA and TC>TT), suggest human APOBEC enzyme activity and previous or ongoing adaption to the human host [24,26]. However, awareness has been raised to not only survey single nucleotide polymorphisms, but rather monitor integrity and stability of the MPXV genomic termini [27]. The double-stranded DNA genome of MPXV contains identical but oppositely oriented sequences (terminal inverted repeats, ITR) of ∼ 6400 bp in length [28]. While essential genes for replication are present in the central, highly conserved core of the genome, the diverse terminal regions of the genome contain genes probably involved in immune evasion and host range [29,30]. Gene gain and loss in the terminal regions are thought to be the main drivers of poxvirus evolution and adaption to the host [31,32]. Deletions in the genomes of clade I MPXV in DRC could be correlated with human-to-human transmission [33], and gene copy number variation has been described as a factor for modulating viral fitness [34]. Furthermore, in VARV, gene loss has been associated with restricted host range, but not severity of disease [35].

Here, we report five MPXV genomes of the multi-country 2022 outbreak, with duplication of several consecutive genes of up to 18,000 bp from the left to the right ITR region as well as from the right to the left ITR region, resulting in gene deletions of up to 17,000 bp in the region of insertion. We could show gene duplication and loss as possible mechanisms of adaption to the human host in the current MPXV outbreak and highlight the need for surveillance of the genome ends rather than or in addition to ongoing monitoring of non-synonymous mutations.

## Methods

So far, 339 samples screened positive for MPXV by qPCR were subjected to Illumina sequencing. DNA was extracted using the QIAamp viral RNA mini kit (Qiagen). Whole genomes shotgun libraries were prepared using the Nextera XT Kit (Illumina). Samples were sequenced on the NextSeq 2000 with approximately 10–30 million reads per sample, using P2 chemistry with 2 × 150 bp. Human background reads were removed, mapping to H. sapiens GRCh38 with bowtie2 v2.3.0. Genomes were constructed with *de novo* assemblies using Spades v3.13.1 and mapping to a reference sequence (Monkeypox/PT0006/2022|sampling_date_20220515, accession ON585033.1). Alignments were prepared with MAFFT V7.307. All genomes in this study were uploaded to NCBI GenBank (MPXV/Germany/2022/RKI335 – MPXV/Germany/2022/RKI339), accession numbers OP696838, OP696839, OP696840, OP696841, OP696842. All sequence data is deposited under the project number PRJEB56807 at the European Nucleotide Archive.

## Results

Whole genome sequencing and *de novo* assembly of 339 genomes of the current multi-country outbreak of MPXV revealed five genomes with duplications, insertions and deletions of a similar pattern (Figure 1, Table 1). Three of the five genomes revealed duplications from the left end to the right end with deletions at the site of insertion. Two of the five genomes showed deletions of genes at the left end, with one genome also having a duplication from the right to the left end.

**Figure 1:**
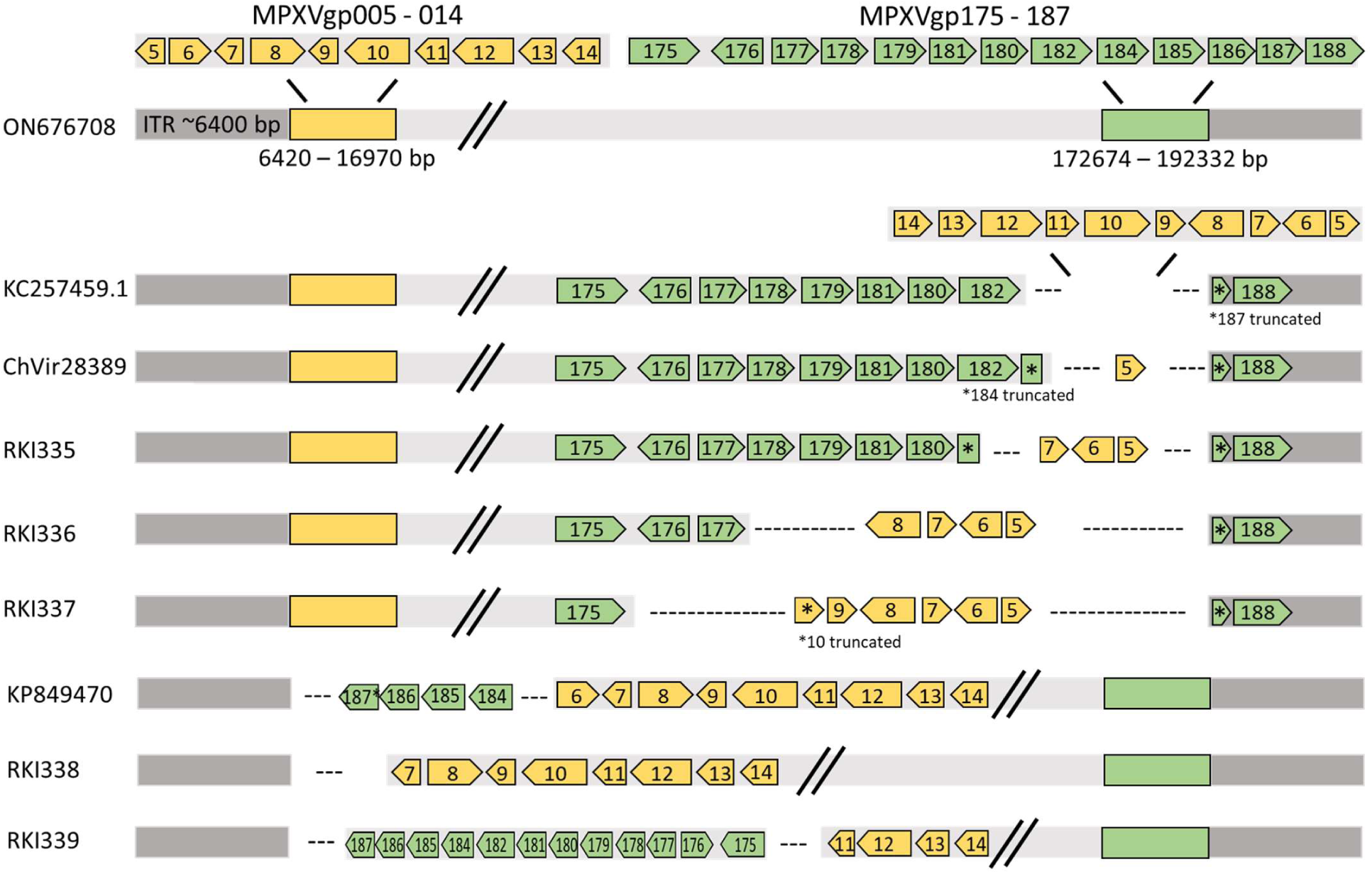
Genes affected by duplications and deletions for left-to-right (KC257459.1, MPXV/Germany/2022/RKI335, MPXV/Germany/2022/RKI336, MPXV/Germany/2022/RKI337, ChVir28389) and right-to-left (KP849470.1 as a representative for 20 genomes of MPXV clade II from 1958–2018) transitions. For left-to-right transitions, the region directly downstream of the left ITR, always including MPXVgp005 and several of the genes MPXVgp006 – MPXVgp014, is copied reverse complementary to upstream of the right ITR, resulting in the extension of the ITR regions. The duplication to the region results in different sizes of deletion, always including a part of MPXVgp187 and the genes MPXVgp175 – MPXVgp186. For right-to-left transitions, the partial MPXV187 and several of the genes MPXVgp175 – MPXVgp186 are copied reverse complementary to the left site of the genome, resulting in increased length of the ITR regions and deletion of the genes MPXVgp005 – MPXVgp010. One of the genomes only shows deletion of the regions MPXV005 and MPXV 006, without right-to-left duplication.

**Table 1:** Duplication and deletion sites and length for left-to-right (KC257459.1, MPXV/Germany/2022/RKI335, MPXV/Germany/2022/RKI336, MPXV/Germany/2022/RKI337, ChVir28389) and right-to-left (KP849470.1 as a representative for 20 genomes of MPXV clade II from 1958–2018) transitions.

|  | Duplication<br>start (bp) | Duplication<br>end (bp) | Deletion<br>start (bp) | Deletion<br>end (bp) | Duplication<br>length (bp) | Deletion<br>length (bp) | Genome<br>length (bp) | ITR length (bp) |
| --- | --- | --- | --- | --- | --- | --- | --- | --- |
| KC257459.1 | 6420 | 16970 | 18644 | 190752 | 10550 | 172108 | 206372 | 17511 |
| MPXV/Germany/2022/RKI335 | 6420 | 9341 | 181859 | 190752 | 2921 | 8893 | 191176 | 9398 |
| MPXV/Germany/2022/RKI336 | 6420 | 10460 | 175449 | 190752 | 4040 | 15303 | 185852 | 10460 |
| MPXV/Germany/2022/RKI337 | 6420 | 11702 | 173826 | 190752 | 5282 | 16926 | 185251 | 11641 |
| ChVir28389 | 6420 | 7278 | 188704 | 190752 | 858 | 2048 | 196039 | 7341 |
| KP849470.1 | 188453 | 190752 | 6420 | 7076 | 2299 | 4777 | 200397 | 9526 |
| MPXV/Germany/2022/RKI338 |  |  | 6420 | 8729 |  | 2309 | 194616 | 6118 |
| MPXV/Germany/2022/RKI339 | 172538 | 190752 | 6420 | 13121 | 18214 | 6701 | 208804 | 24695 |

For left-to-right duplications, the length of the duplications could be correlated with increased length of deletions at the insertion site, increased length of ITR regions.

Genome MPXV/Germany/2022/RKI335 showed a duplication from the region downstream of the left ITR, including the genes MPXVgp005–007, to the regions upstream of the right ITR between genes MPXVgp182 and MPXVgp188, resulting in truncation of MPXVgp184 and MPXVgp187 and complete deletion of MPXVgp185–186. The size of duplication (2921 bp) to the right end and the deletion of 8893 bp resulted in a total genome length of 191,176 bp (ON676708: 197,173 bp) and the extension of the ITR regions from ∼6,400 bp to 9,398 bp. Similar duplications and deletions were shown for MPXV/Germany/2022/RKI336 with a duplication of the genes MPXVgp005–008 and insertion at the right genome end between MPXVgp177 and MPXVgp188, with truncation of MPXVgp187 and complete deletion of genes MPXVgp178–186. The duplication of 4,040 bp and deletion of 15,303 bp resulted in a genome length of 185,852 bp with ITRs of 10,460 bp. Genome 3 (MPXV/Germany/2022/RKI337) showed the same pattern of duplication and deletion, with the largest duplication of 5,282 bp (MPXVgp005 – MPXVgp010 [truncated]) to the right end with also the largest deletion of 16,926 bp (MPXVgp176 – MPXVgp187 [truncated]). The genome length of MPXV/Germany/2022/RKI337 was 185,251 bp with ITRs of 11,641 bp.

Performing whole genome alignments with all of the MPXV genomes available at NCBI GenBank, such duplications from left to right were also found in a clade I MPXV genome from 2005 (KC257459.1, Monkeypox strain Sudan 2005) and in a genome from the recent outbreak (hMPXV/Germany/BE-ChVir28389/2022|EPI_ISL_13890482|2022-06-08) recently published [36,37]. The MPXV genome from the Sudan showed a duplication of 10,550 bp from the same region after the left ITR to the right ITR, with a deletion of 2,108 bp. The recently described duplication in genome E-ChVir28389 showed a duplication of 858 bp from the left to the right ITR, with a deletion of 2,048 bp.

Duplications from the right to the left end were also revealed, with MPXV/Germany/2022/RKI339 having the largest duplication of 18,214 (genes MPXVgp174 [truncated] – MPXVgp187 [truncated]) to the left end, resulting in the deletion of genes MPXVgp005 – MPXVgp010 (6,701 bp). The genome length was increased to 208,804 bp with extended ITRs of 24,695 bp. Genome MPXV/Germany/2022/RKI338 showed a deletion of 2,309 bp in the same region downstream of the left ITR, but without an insertion. Similar duplications were noticed in 20 genomes of clade II MPXV from 1958 to 2018.

## Discussion

During the SARS-CoV-2 pandemic, genomic surveillance has been focusing on single non-synonymous mutations, arising frequently and resulting in different phenotypes and variants and affecting virulence and host escape [38]. Poxviruses evolve much slower than RNA viruses, and it is rather hypothesized that all of today’s OPVs descended from a common ancestor and evolved by loss of accessory genes, including those relevant for host adaption [39]. Deleted genes could be correlated with loss of an antigenic signal or continued evolutionary persistence by the attenuation of disease [40]. In contrast, acquired genes may support evasion of host defenses. For example, “genome accordions” have been described as a rapid mechanism for adaption of Vaccinia virus (VACV), including amplification of genes under selective pressure, accumulation of point mutations and the following deletion of the additional genes [34]. CPXV has the highest number of accessory genes and the broadest host range, while VARV has the smallest set of such genes and no host other than humans [41]. VARV strains from human archeological remains from the Viking Age were possibly widespread, but less pathogenic, supported by the discovery of accessory genes which are lost in modern-day VARV [35]. It is hypothesized that the variola-like ancestors’ host reservoir were rodents, as described for many modern OPVs today, including MPXV, and evolved into modern highly pathogenic VARV after spill-over and adaption to the human host [42]. Gene gain and loss have been described previously for MPXV. In 60 samples from humans in DRC, collected between 2005 and 2007, 10 MPXV genomes showed a 625-bp deletion directly upstream of the right ITR, as shown for the left-to-right transitions in our study, although no gene insertion could be observed. The deletion included one full and one truncated gene and seemed to correlate with human-to-human transmission [33]. Furthermore, as shown in Figure 1, 20 clade II genomes obtained from GenBank contained duplications from right to left. Most of the genomes were derived from chimpanzees sampled in 2017 and rodents from the US outbreak in 2003 [43,44]. The Sudan clade I genome with the largest insertion, KC257459.1, was sampled from a human MPXV outbreak in the Southern Sudan in 2005 [37]. Although sequences were obtained from primary clinical material directly, it is mentioned that the insertion was partially lost after the second passage in cell culture.

We could show similar left-to-right and right-to-left duplications, resulting in large deletions in five genomes of the 2022 human MPXV outbreak. All left-to-right duplications included one or more of the genes MPXVgp005 – MPXV014 with deletion of one or more of the genes MPXVgp175 – MPXVgp187, or the other way around for right-to-left duplications. For all left-to-right duplications, at least genes MPXVgp005 – MPXVgp007 were included. Hypothesized protein functions are shown in supplementary table 1. MPXVgp005 is not homologous to any gene of the OPV genus and its function is unknown. Furthermore, after reannotation, the fragmented gene had been removed [45]. MPXVgp006 is an epidermal growth factor-like protein, homologous to the C11R Vaccinia virus gene. It was shown that deletion of C11R reduced NF-κB activation in cell culture [46]. It has been discussed whether MPXVgp007 has been annotated correctly [45]. It has no homologue in any other OPV. Further duplicated genes are MPXVgp008 (zinc-finger protein, inhibition of apoptosis), MPXVgp009 (functional inhibitor of interleukin 18) and MPXVgp010 (truncated ankyrin/host range protein). Furthermore, all left-to-right duplications led to the deletion of at least MPXVgp184 (truncated) – MPXVgp187 (truncated). Again, MPXVgp184 and MPXVgp186 have no OPV homologues and are discussed to be annotated incorrectly [45]. MPXVgp185 is homologous to Vaccinia B22R, an immune-modulating gene with homology to serine protease inhibitors [47]. Deletion of B22R has been shown to decrease viral replication and virulence [48]. For MPXV/Germany/2022/RKI336 and MPXV/Germany/2022/RKI337 genes, MPXVgp180 (B19R) and MPXV182 (B21R) are further deleted – being two out of the six candidates identified as main possible virulence genes in MPXV [45]. Another one of these candidates, D10L (MPXVgp013), together with B19R and B21R, is duplicated in MPXV/Germany/2022/RKI339, and a fourth candidate is duplicated in the Sudan KC257459 genome shown in the results.

One may speculate whether such genome rearrangements lead to different MPXV phenotypes and are a sign of ongoing adaption to the human host. Relevant accessory host range genes are duplicated in the case of right-to-left duplications, but are also deleted in the case of left-to-right duplications. Furthermore, the function of genes is often hypothesized or adopted from VACV or CPXV. Duplications near the termini have been described frequently for CPXV grown in chicken embryos, induced by crossover recombinational events between two genomes in opposite direction [49]. However, the large insertions are described to be unstable, being lost after passages in cell culture [50]. It is therefore possible that the genomes in this study are intermediates of ongoing evolution, finally leading to reduced accessory genes in the genome ends as adaption to the human host, as could have happened for genome MPXV/Germany/2022/RKI338 only showing the deletion after possible loss of the insertion, and it has been described for VARV [35]. Another famous example is VACV which was used as a vaccine in the eradication of smallpox. VACV is thought to be derived from what today is known as horsepox virus, which has possibly been used by Edward Jenner when inventing vaccination [51,52]. One of the main differences to today’s VACV are 10.7-kb and 5.5-kb insertions at the genomes’ ends. Vaccines from the US Civil War era (1859–1873) consist of true horsepox virus including the insertions, but also VACV-like viruses with and without the insertions as well as intermediates with only one insertion and end-to-end duplications, including a right-to-left duplication covering the MPXV homologue genes MPXVgp172 – MPXVgp182 [53,54]. Early VACV was disseminated through serial arm-to-arm transmission, with possible gene loss as an adaption to the host. The origin and natural host of the ancestor of VACV is still a mystery, but today’s VACV have only caused sporadic outbreaks and were not able to re-emerge in a natural host [55].

In this study, we only found five out of 339 MPXV genomes (1.47 %) with rearrangements. Ongoing sequencing efforts will have to show whether some of the described insertions and deletions will become fixed as a possible adaption to the new human host, or whether they are neutral or even disadvantageous for the virus. The MPXV of the 2022 outbreak may have become adapted to the human host, as continuous human-to-human transmission is seen for the first time since the discovery of MPXV in 1958. In addition, the mode of transmission and the phenotype of disease has changed [56,57]. Although adaption to the host is not necessarily correlated with increased pathogenicity and virulence, during the ongoing pandemic it is not known how the virus will evolve in future. Intensified vaccination and changes in human behavior could exert selective pressure on the virus, with mutations, insertions and deletions being advantageous for transmission or immune escape. Although MPXV daily cases worldwide are decreasing (as of September 27, 2022), it is likely that MPXV outbreaks will continue to appear, with different strains circulating, some of which seemingly adapted to the human host [58].

## Conclusion

Rearrangements, including gene transitions to the opposite genome end and deletions at the insertion site of the genomes of the MPXV 2022 multi-country outbreak, can be monitored by whole genome sequencing. Gene gain and loss at the terminal ends of the MPXV genome have been described as an adaption to the host.

## Acknowledgments

We kindly thank Ursula Erikli for copy-editing and Ute Kramer for technical assistance.

## Ethics statement

The studies involving human participants were reviewed and approved by the Ärztekammer Berlin (Berlin Medical Association; #Eth-44/22). The patients provided their written informed consent to sequencing.

## Conflict of interest

None declared.

## Authors’ contributions

Annika Brinkmann designed the assay, analyzed the data and wrote the manuscript. Katharina Pape analyzed the samples and the data. Andrea Thürmer performed Illumina sequencing of the samples. Janine Michel and Claudia Kohl and Daniel Bourquain analyzed the suspected MPXV specimens. Lars Schaade and Andreas Nitsche conceptualized the approach and wrote the manuscript.

## Notes

### Competing Interest Statement

The authors have declared no competing interest.

## References

1. World Health Organization. Global Commission for the Certification of Smallpox Eradication. The achievement of global eradication of smallpox: final report of the Global Commission for the Certification of Smallpox Eradication, Geneva, December 1979. Geneva: WHO, 1979.

2. Lloyd-Smith, J.O. Vacated niches, competitive release and the community ecology of pathogen eradication. Philos Trans R Soc Lond B Biol Sci 2013, 368, 20120150, doi:10.1098/rstb.2012.0150.

3. Reynolds, M.G.; Carroll, D.S.; Karem, K.L. Factors affecting the likelihood of monkeypox’s emergence and spread in the post-smallpox era. Curr Opin Virol 2012, 2, 335–343, doi:10.1016/j.coviro.2012.02.004.

4. Ladnyj, I.D.; Ziegler, P.; Kima, E. A human infection caused by monkeypox virus in Basankusu Territory, Democratic Republic of the Congo. Bull World Health Organ 1972, 46, 593–597.

5. Centers for Disease Control and Prevention (CDC). Multistate outbreak of monkeypox--Illinois, Indiana, and Wisconsin, 2003. MMWR. Morb Mortal Wkly Rep 2003, 52, 537–540.

6. Formenty, P.; Muntasir, M.O.; Damon, I.; Chowdhary, V.; Opoka, M.L.; Monimart, C.; Mutasim, E.M.; Manuguerra, J.C.; Davidson, W.B.; Karem, K.L.; et al. Human monkeypox outbreak caused by novel virus belonging to Congo Basin clade, Sudan, 2005. Emerg Infect Dis 2010, 16, 1539–1545, doi:10.3201/eid1610.100713.

7. Yinka-Ogunleye, A.; Aruna, O.; Dalhat, M.; Ogoina, D.; McCollum, A.; Disu, Y.; Mamadu, I.; Akinpelu, A.; Ahmad, A.; Burga, J.; et al. Outbreak of human monkeypox in Nigeria in 2017–18: a clinical and epidemiological report. Lancet Infect Dis 2019, 19, 872–879, doi:10.1016/s1473-3099(19)30294-4.

8. Rezza, G. Emergence of human monkeypox in west Africa. Lancet Infect Dis 2019, 19, 797–799, doi:10.1016/s1473-3099(19)30281-6.

9. Erez, N.; Achdout, H.; Milrot, E.; Schwartz, Y.; Wiener-Well, Y.; Paran, N.; Politi, B.; Tamir, H.; Israely, T.; Weiss, S.; et al. Diagnosis of Imported Monkeypox, Israel, 2018. Emerg Infect Dis 2019, 25, 980–983, doi:10.3201/eid2505.190076.

10. Yong, S.E.F.; Ng, O.T.; Ho, Z.J.M.; Mak, T.M.; Marimuthu, K.; Vasoo, S.; Yeo, T.W.; Ng, Y.K.; Cui, L.; Ferdous, Z.; et al. Imported Monkeypox, Singapore. Emerg Infect Dis 2020, 26, 1826–1830, doi:10.3201/eid2608.191387.

11. Vaughan, A.; Aarons, E.; Astbury, J.; Brooks, T.; Chand, M.; Flegg, P.; Hardman, A.; Harper, N.; Jarvis, R.; Mawdsley, S.; et al. Human-to-Human Transmission of Monkeypox Virus, United Kingdom, October 2018. Emerg Infect Dis 2020, 26, 782–785, doi:10.3201/eid2604.191164.

12. Durski, K.N.; McCollum, A.M.; Nakazawa, Y.; Petersen, B.W.; Reynolds, M.G.; Briand, S.; Djingarey, M.H.; Olson, V.A.; Damon, I.K.; Khalakdina, A. Emergence of Monkeypox — West and Central Africa, 1970–2017. MMWR. Morb Mort Wkly Rep 2018, 67, 306–310, doi:10.15585/mmwr.mm6710a5.

13. Learned, L.A.; Reynolds, M.G.; Wassa, D.W.; Li, Y.; Olson, V.A.; Karem, K.; Stempora, L.L.; Braden, Z.H.; Kline, R.; Likos, A.; et al. Extended interhuman transmission of monkeypox in a hospital community in the Republic of the Congo, 2003. Am JTrop Med Hyg 2005, 73, 428–434, doi:10.4269/ajtmh.2005.73.428.

14. Jezek, Z.; Arita, I.; Mutombo, M.; Dunn, C.; Nakano, J.H.; Szczeniowski, M. Four generations of probable person-to-person transmission of human monkeypox. Am J Epidemiol 1986, 123, 1004–1012, doi:10.1093/oxfordjournals.aje.a114328.

15. Nolen, L.D.; Osadebe, L.; Katomba, J.; Likofata, J.; Mukadi, D.; Monroe, B.; Doty, J.; Hughes, C.M.; Kabamba, J.; Malekani, J.; et al. Extended Human-to-Human Transmission during a Monkeypox Outbreak in the Democratic Republic of the Congo. Emerg Infect Dis 2016, 22, 1014–1021, doi:10.3201/eid2206.150579.

16. Khodakevich, L.; Jezek, Z.; Kinzanzka, K. Isolation of monkeypox virus from wild squirrel infected in nature. Lancet 1986, 1, 98–99, doi:10.1016/s0140-6736(86)90748-8.

17. Hutson, C.L.; Nakazawa, Y.J.; Self, J.; Olson, V.A.; Regnery, R.L.; Braden, Z.; Weiss, S.; Malekani, J.; Jackson, E.; Tate, M.; et al. Laboratory Investigations of African Pouched Rats (Cricetomys gambianus) as a Potential Reservoir Host Species for Monkeypox Virus. PLOS Negl Trop Dis 2015, 9, e0004013, doi:10.1371/journal.pntd.0004013.

18. Parker, S.; Buller, R.M. A review of experimental and natural infections of animals with monkeypox virus between 1958 and 2012. Future Virol 2013, 8, 129–157, doi:10.2217/fvl.12.130.

19. Mahase, E. Seven monkeypox cases are confirmed in England. BMJ 2022, 377, o1239, doi:10.1136/bmj.o1239.

20. Centers for Disease Control and Prevention. CDC and Health Partners Responding to Monkeypox Case in the U.S. https://www.cdc.gov/media/releases/2022/s0518-monkeypox-case.html Accessed 2022 Oct 12.

21. Reuters. Portugal identifies five monkeypox infections, Spain has 23 suspected cases. https://www.reuters.com/business/healthcare-pharmaceuticals/portugal-identifies-five-monkeypox-infections-spain-has-23-suspected-cases-2022-05-18/ Accessed 2022 Oct 12.

22. Centers for Disease Control and Prevention. 2022 Monkeypox Outbreak Global Map. https://www.cdc.gov/poxvirus/monkeypox/response/2022/world-map.html Accessed 2022 Oct 12.

23. Likos, A.M.; Sammons, S.A.; Olson, V.A.; Frace, A.M.; Li, Y.; Olsen-Rasmussen, M.; Davidson, W.; Galloway, R.; Khristova, M.L.; Reynolds, M.G.; et al. A tale of two clades: monkeypox viruses. J Gen Virol 2005, 86, 2661–2672, doi:10.1099/vir.0.81215-0.

24. Isidro, J.; Borges, V.; Pinto, M.; Sobral, D.; Santos, J.D.; Nunes, A.; Mixao, V.; Ferreira, R.; Santos, D.; Duarte, S.; et al. Phylogenomic characterization and signs of microevolution in the 2022 multi-country outbreak of monkeypox virus. Nat Med 2022, 28, 1569–1572, doi:10.1038/s41591-022-01907-y.

25. Gigante, C.M.; Korber, B.; Seabolt, M.H.; Wilkins, K.; Davidson, W.; Rao, A.K.; Zhao, H.; Hughes, C.M.; Minhaj, F.; Waltenburg, M.A.; et al. Multiple lineages of Monkeypox virus, detected in the United States, 2021-2022. bioRxiv 2022, 2022.06.10.495526, doi:10.1101/2022.06.10.495526.

26. Firth, C.; Kitchen, A.; Shapiro, B.; Suchard, M.A.; Holmes, E.C.; Rambaut, A. Using time-structured data to estimate evolutionary rates of double-stranded DNA viruses. Mol Biol Evol 2010, 27, 2038–2051, doi:10.1093/molbev/msq088.

27. Pfaff, F.; Hoffmann, D.; Beer, M. Monkeypox genomic surveillance will challenge lessons learned from SARS-CoV-2. Lancet 2022, 400, 22–23, doi:10.1016/s0140-6736(22)01106-0.

28. Shchelkunov, S.N.; Totmenin, A.V.; Safronov, P.F.; Mikheev, M.V.; Gutorov, V.V.; Ryazankina, O.I.; Petrov, N.A.; Babkin, I.V.; Uvarova, E.A.; Sandakhchiev, L.S.; et al. Analysis of the monkeypox virus genome. Virology 2002, 297, 172–194, doi:10.1006/viro.2002.1446.

29. Alcami, A.; Koszinowski, U.H. Viral mechanisms of immune evasion. Trends Microbiol 2000, 8, 410–418, doi:10.1016/S0966-842X(00)01830-8.

30. Moss, B.; Shisler, J.L. Immunology 101 at poxvirus U: Immune evasion genes. Semin Immunol 2001, 13, 59–66, doi:10.1006/smim.2000.0296.

31. Esteban, D.J.; Hutchinson, A.P. Genes in the terminal regions of orthopoxvirus genomes experience adaptive molecular evolution. BMC Genomics 2011, 12, 261, doi:10.1186/1471-2164-12-261.

32. Hughes, A.L.; Friedman, R. Poxvirus genome evolution by gene gain and loss. Mol Phylogenet Evol 2005, 35, 186–195, doi:10.1016/j.ympev.2004.12.008.

33. Kugelman, J.R.; Johnston, S.C.; Mulembakani, P.M.; Kisalu, N.; Lee, M.S.; Koroleva, G.; McCarthy, S.E.; Gestole, M.C.; Wolfe, N.D.; Fair, J.N.; et al. Genomic variability of monkeypox virus among humans, Democratic Republic of the Congo. Emerg Infect Dis 2014, 20, 232–239, doi:10.3201/eid2002.130118.

34. Elde, N.C.; Child, S.J.; Eickbush, M.T.; Kitzman, J.O.; Rogers, K.S.; Shendure, J.; Geballe, A.P.; Malik, H.S. Poxviruses deploy genomic accordions to adapt rapidly against host antiviral defenses. Cell 2012, 150, 831–841, doi:10.1016/j.cell.2012.05.049.

35. Mühlemann, B.; Vinner, L.; Margaryan, A.; Wilhelmson, H.; de la Fuente Castro, C.; Allentoft, M.E.; de Barros Damgaard, P.; Hansen, A.J.; Holtsmark Nielsen, S.; Strand, L.M.; et al. Diverse variola virus (smallpox) strains were widespread in northern Europe in the Viking Age. Science 2020, 369, eaaw8977, doi:10.1126/science.aaw8977.

36. Jones, T.C.; Schneider, J.; Mühlemann, B.; Veith, T.; Beheim-Schwarzbach, J.; Tesch, J.; Schmidt, M.L.; Walper, F.; Bleicker, T.; Isner, C.; et al. Genetic variability, including gene duplication and deletion, in early sequences from the 2022 European monkeypox outbreak. bioRxiv 2022, 2022.07.23.501239, doi:10.1101/2022.07.23.501239.

37. Nakazawa, Y.; Emerson, G.L.; Carroll, D.S.; Zhao, H.; Li, Y.; Reynolds, M.G.; Karem, K.L.; Olson, V.A.; Lash, R.R.; Davidson, W.B.; et al. Phylogenetic and ecologic perspectives of a monkeypox outbreak, southern Sudan, 2005. Emerg Infect Dis 2013, 19, 237–245, doi:10.3201/eid1902.121220.

38. Oude Munnink, B.B.; Worp, N.; Nieuwenhuijse, D.F.; Sikkema, R.S.; Haagmans, B.; Fouchier, R.A.M.; Koopmans, M. The next phase of SARS-CoV-2 surveillance: real-time molecular epidemiology. Nat Med 2021, 27, 1518–1524, doi:10.1038/s41591-021-01472-w.

39. Senkevich, T.G.; Yutin, N.; Wolf, Y.I.; Koonin, E.V.; Moss, B. Ancient Gene Capture and Recent Gene Loss Shape the Evolution of Orthopoxvirus-Host Interaction Genes. mBio 2021, 12, e0149521, doi:10.1128/mBio.01495-21.

40. McLysaght, A.; Baldi, P.F.; Gaut, B.S. Extensive gene gain associated with adaptive evolution of poxviruses. Proc Natl Acad Sci U S A 2003, 100, 15655–15660, doi:10.1073/pnas.2136653100.

41. Lefkowitz, E.J.; Wang, C.; Upton, C. Poxviruses: past, present and future. Virus Res 2006, 117, 105–118, doi:10.1016/j.virusres.2006.01.016.

42. Alcamí, A. Was smallpox a widespread mild disease? Science 2020, 369, 376–377, doi:10.1126/science.abd1214.

43. Patrono, L.V.; Pleh, K.; Samuni, L.; Ulrich, M.; Rothemeier, C.; Sachse, A.; Muschter, S.; Nitsche, A.; Couacy-Hymann, E.; Boesch, C.; et al. Monkeypox virus emergence in wild chimpanzees reveals distinct clinical outcomes and viral diversity. Nat Microbiol 2020, 5, 955–965, doi:10.1038/s41564-020-0706-0.

44. Mauldin, M.R.; McCollum, A.M.; Nakazawa, Y.J.; Mandra, A.; Whitehouse, E.R.; Davidson, W.; Zhao, H.; Gao, J.; Li, Y.; Doty, J.; et al. Exportation of Monkeypox Virus From the African Continent. J Infect Dis 2022, 225, 1367–1376, doi:10.1093/infdis/jiaa559.

45. Chen, N.; Li, G.; Liszewski, M.K.; Atkinson, J.P.; Jahrling, P.B.; Feng, Z.; Schriewer, J.; Buck, C.; Wang, C.; Lefkowitz, E.J.; et al. Virulence differences between monkeypox virus isolates from West Africa and the Congo basin. Virology 2005, 340, 46–63, doi:10.1016/j.virol.2005.05.030.

46. Martin, S.; Harris, D.T.; Shisler, J. The C11R gene, which encodes the vaccinia virus growth factor, is partially responsible for MVA-induced NF-kappaB and ERK2 activation. J Virol 2012, 86, 9629–9639, doi:10.1128/JVI.06279-11.

47. Legrand, F.A.; Verardi, P.H.; Chan, K.S.; Peng, Y.; Jones, L.A.; Yilma, T.D. Vaccinia viruses with a serpin gene deletion and expressing IFN-gamma induce potent immune responses without detectable replication in vivo. Proc Natl Acad Sci U S A 2005, 102, 2940–2945, doi:10.1073/pnas.0409846102.

48. Legrand, F.A.; Verardi, P.H.; Jones, L.A.; Chan, K.S.; Peng, Y.; Yilma, T.D. Induction of potent humoral and cell-mediated immune responses by attenuated vaccinia virus vectors with deleted serpin genes. J Virol 2004, 78, 2770–2779, doi:10.1128/jvi.78.6.2770-2779.2004.

49. Pickup, D.J.; Ink, B.S.; Parsons, B.L.; Hu, W.; Joklik, W.K. Spontaneous deletions and duplications of sequences in the genome of cowpox virus. Proc Natl Acad Sci U S A 1984, 81, 6817–6821, doi:10.1073/pnas.81.21.6817.

50. Vallée, G.; Norris, P.; Paszkowski, P.; Noyce, R.S.; Evans, D.H. Vaccinia Virus Gene Acquisition through Nonhomologous Recombination. J Virol 2021, 95, e0031821, doi:10.1128/JVI.00318-21.

51. Schrick, L.; Tausch, S.H.; Dabrowski, P.W.; Damaso, C.R.; Esparza, J.; Nitsche, A. An Early American Smallpox Vaccine Based on Horsepox. N Engl J Med 2017, 377, 1491–1492, doi:10.1056/NEJMc1707600.

52. Esparza, J.; Nitsche, A.; Damaso, C.R. Beyond the myths: Novel findings for old paradigms in the history of the smallpox vaccine. PLoS Pathog 2018, 14, e1007082, doi:10.1371/journal.ppat.1007082.

53. Duggan, A.T.; Klunk, J.; Porter, A.F.; Dhody, A.N.; Hicks, R.; Smith, G.L.; Humphreys, M.; McCollum, A.M.; Davidson, W.B.; Wilkins, K.; et al. The origins and genomic diversity of American Civil War Era smallpox vaccine strains. Genome Biol 2020, 21, 175, doi:10.1186/s13059-020-02079-z.

54. Brinkmann, A.; Souza, A.R.V.; Esparza, J.; Nitsche, A.; Damaso, C.R. Re-assembly of nineteenth-century smallpox vaccine genomes reveals the contemporaneous use of horsepox and horsepox-related viruses in the USA. Genome Biol 2020, 21, 286, doi:10.1186/s13059-020-02202-0.

55. Abrahao, J.S.; Campos, R.K.; Trindade Gde, S.; Guimaraes da Fonseca, F.; Ferreira, P.C.; Kroon, E.G. Outbreak of severe zoonotic vaccinia virus infection, Southeastern Brazil. Emerg Infect Dis 2015, 21, 695–698, doi:10.3201/eid2104.140351.

56. Sah, R.; Abdelaal, A.; Reda, A.; Katamesh, B.E.; Manirambona, E.; Abdelmonem, H.; Mehta, R.; Rabaan, A.A.; Alhumaid, S.; Alfouzan, W.A.; et al. Monkeypox and Its Possible Sexual Transmission: Where Are We Now with Its Evidence? Pathogens 2022, 11, 924, doi:10.3390/pathogens11080924.

57. Davido, B.; D’Anglejan, E.; Jourdan, J.; Robinault, A.; Davido, G. Monkeypox 2022 outbreak: cases with exclusive genital lesions. J Travel Med 2022, 29, taac077, doi:10.1093/jtm/taac077.

58. World Health Organization. 2022 Monkeypox Outbreak: Global Trends. https://worldhealthorg.shinyapps.io/mpx_global/ Accessed 2022 Oct 17.

